# Genomic resources and toolkits for developmental study of whip spiders (Amblypygi) provide insights into arachnid genome evolution and antenniform leg patterning

**DOI:** 10.1101/2020.07.03.185769

**Authors:** Guilherme Gainett, Prashant P. Sharma

## Abstract

**Background:** The resurgence of interest in the comparative developmental study of chelicerates has led to important insights, such as the discovery of a genome duplication shared by spiders and scorpions, inferred to have occurred at the most recent common ancestor of Arachnopulmonata (a clade comprised of the five book lung-bearing arachnid orders). Nonetheless, several arachnid groups remain understudied in the context of development and genomics, such as the order Amblypygi (whip spiders). The phylogenetic position of Amblypygi in Arachnopulmonata posits them as an interesting group to test the incidence of the proposed genome duplication in the common ancestor of Arachnopulmonata, as well as the degree of retention of duplicates over 450 Myr. Moreover, whip spiders have their first pair of walking legs elongated and modified into sensory appendages (a convergence with the antenna of mandibulates), but the genetic patterning of these antenniform legs has never been investigated.

**Results:** We established genomic resources and protocols for cultivation of embryos and gene expression assays by in situ hybridization to study the development of the whip spider *Phrynus marginemaculatus*. Using embryonic transcriptomes from three species of Amblypygi, we show that the ancestral whip spider exhibited duplications of all ten Hox genes. We deploy these resources to show that paralogs of the leg gap genes *dachshund* and *homothorax* retain arachnopulmonate-specific expression patterns in *P. marginemaculatus*. We characterize the expression of leg gap genes *Distal-less, dachshund-1/2* and *homothorax-1/2* in the embryonic antenniform leg and other appendages, and provide evidence that allometry, and by extension the antenniform leg fate, is specified early in embryogenesis.

**Conclusion:** This study is the first step in establishing *P. marginemaculatus* as a model for modern chelicerate evolutionary developmental study, and provides the first resources sampling whip spiders for comparative genomics. Our results suggest that Amblypygi share a genome duplication with spiders and scorpions, and set up a framework to study the genetic specification of antenniform legs. Future efforts to study whip spider development must emphasize the development of tools for functional experiments in *P. marginemaculatus*.

## Background

In the past 25 years, comparative developmental study of chelicerates has resurged with the integration of molecular biology and genomics, spearheaded by developmental genetic investigations of the spiders *Cupiennius salei* and *Parasteatoda tepidariorum* [1-6]. Such works have revealed several important aspects of chelicerate development, such as the molecular mechanism of dorso-ventral axis patterning [3,7-9], segmentation of the prosoma [1,10-13] and opisthosoma [2,13], and specification of prosomal versus opisthosomal fate [14,15]. A few non-spider chelicerate models have also emerged in recent years and provided new perspectives on chelicerate development, such as the horseshoe crab *Limulus polyphemus* (Xiphosura) [16,17], the mites *Archegozetes longisetosus* [18-21] and *Tetranychus urticae* (Acariformes) [22-24], the tick *Rhipicephalus microplus* (Parasitiformes) [25], the Arizona bark scorpion *Centruroides sculpturatus* (Scorpiones) [26] and the harvestmen *Phalangium opilio* (Opiliones) [27-30]. Studies in *A. longisetosus* and *C. sculpturatus* have suggested that changes in Hox gene number and expression domains are responsible for such phenomena as the reduced segmentation of mites and the supernumerary posterior appendage identities of scorpions [18,26].

In addition to the insights into chelicerate development, one of the major recent outcomes of increasing availability of genomic resources in in the group was the discovery of a whole or partial genome duplication in spiders and scorpions. This genome duplication is inferred to trace back to the most recent common ancestor of the recently proposed clade Arachnopulmonata, which includes spiders, scorpions, and three other arachnid orders with book lungs [6,26,31-33]. This duplication event is also inferred to be independent from the multiple rounds of whole genome duplication undergone by horseshoe crabs (Xiphosura) [34,35]. The duplication in spiders and scorpions encompasses several important developmental genes, such as the homeobox family [6,32]. Excitingly, there is increasing evidence of divergent expression and function of paralogs, such as in the case of Hox genes, leg gap genes and Retinal Determination Gene Network homologs [6,26,36-40].

Nevertheless, knowledge on the development of several arachnid orders still remains scarce or entirely unexplored in the contexts of embryological study and genomic architecture. One particularly interesting arachnid order whose genomic evolution and developmental biology remains largely unexplored is Amblypygi. Commonly known as whip spiders, Amblypygi comprise approximately 220 described species of nocturnal predators that are distributed primarily in the tropics and subtropics worldwide [41,42]. Amblypygi is part of the clade Pedipalpi (together with the orders Thelyphonida and Schizomida), which in turn is sister group to spiders (Araneae) [31,43-45]. Considering the phylogenetic position of Amblypygi within Arachnopulmonata, a better understanding of Amblypygi genomics is anticipated to facilitate exploration of the evolutionary outcomes of the genome duplication in Arachnopulmonata, and better characterizing the extent to which arachnopulmonate orders retained and specialized the ensuing paralogous genes. Unfortunately, the embryology of the order Amblypygi is known only from the works of Pereyaslawzewa [46], a brief mention in Strubell [47] (“Phrynidae”), Gough [48], and the comprehensive study of Weygoldt [49] in *Phrynus marginemaculatus* (formerly *Tarantula marginemaculata*).

Beyond the interest in their genomic architecture, whip spiders possess a fascinating biology and natural history. Whip spiders are emerging as model organisms in behavioral ecology, learning, and neurophysiology in arachnids [50,51]. Their pedipalps, the second pair of appendages, are robust raptorial devices used for striking prey, including mostly arthropods, and even small vertebrates [41,50,52]. However, their most conspicuous characteristic and namesake is their “whip”, a modified antenniform first walking leg that is not used for locomotion, but instead as a sensory appendage. The antenniform legs are extremely elongated relative to the other leg pairs, and the tibia and tarsus are pseudo-segmented into hundreds of articles that harbor an array of chemo-, thermo-, hygro- and mechanoreceptive sensilla [53-55]. The peripheral circuitry of the antenniform legs is complex, exhibiting peripheral synapses and giant neurons which are possibly involved in fast sensory responses [55-57]. The antenniform legs also have been shown to be important for foraging, mating, and intrasexual contests [58,59]. Notably, concentration of sensory structures, elongation and pseudo-segmentation of these legs constitute a striking convergence with the antenna of mandibulate arthropods (Pancrustacea and Myriapoda). Evidence from Hox gene expression and functional experiments support the view that the antenna of mandibulates is positionally homologous to the chelicera of chelicerates (deutocerebral appendage) [21,30]. Despite their different positions along the antero-posterior axis of the body, the serial homology of antennae (mandibulates) and antenniform legs (whip spiders) to walking legs constitutes a potentially useful comparison to address whether a striking morphological convergence is associated with convergence in mechanisms of genetic patterning.

Most of the knowledge about the genetics of antenna versus leg fate specification in arthropods was discovered in the fruit fly *Drosophila melanogaster* and involves the regulation by leg gap genes and Hox genes. Broadly overlapping domains of leg gap genes *homothorax* (*hth*), *dachshund* (*dac*) and *Distal-less* (*Dll*) in the antenna promote antennal fate through activation of downstream antennal determinants such as *spineless* [60-63] (reviewed by [64]); leg identity in thoracic appendages is mediated by mutually antagonistic *hth* (proximal), *dac* (median) and *Dll* (distal) domains, and repression of *hth* by the Hox gene *Antennapedia* (*Antp*) [62,65,66]. Ectopic expression of *hth* on the leg discs results in leg-to-antenna transformation, whereas ectopic *Antp* in the antennal disc results in antenna-to-leg transformation [62,65,66]. In chelicerates, similar to *D. melanogaster*, the chelicera (antennal positional homolog) has broadly overlapping domains of *Dll* and *hth*; legs and pedipalps lack a distal *hth*, and a proximal *Dll* expression domain, indicating a conservation of expression of positional homologs [28,67-69]. In line with this evidence, knockdown of *hth* in the harvestmen *Phalangium opilio* results in chelicera-to-leg transformation (and also partial pedipalp-to-leg transformation) [30]. In contrast to *D. melanogaster* and other insects, arachnid *Antp* is not expressed in the leg-bearing segments, and knockdown of this gene in the spider *Parasteatoda tepidariorum* has revealed a role in repressing limbs in the opisthosoma [14]. Expression of *dac* in a mid-domain of chelicerate pedipalps and legs suggests a conserved role in mid segment patterning, and, accordingly, knockdown of *dac* in the harvestmen *P. opilio* and the spider *P. tepidariorum* results in defects on the mid-leg segments [27,38]. Additionally, loss of a *dac* domain of the chelicera of spiders in comparison to harvestman has been implicated in the transition from three-to two-segmented chelicera [27,28].

Towards revitalizing embryological studies in Amblypygi, we generated herein protocols and comprehensive genomic resources for developmental genetic study of the whip spider *Phrynus marginemaculatus* (Fig. 1A–B), resuming the seminal studies begun by Peter Weygoldt with this promising model species. Leveraging a comprehensive developmental transcriptome for this species, we first investigated the presence and retention of systemic gene duplications in Amblypygi predicted from their phylogenetic position in Arachnopulmonata; to mitigate biases stemming from survey of a data point from a single exemplar, we included as well the embryonic transcriptomes we recently generated for two species of the distantly related family Charinidae (*Charinus ioanniticus* and *Charinus israelensis*) [40,41,70]. As a first step towards elucidating the patterning of antenniform legs, we investigated a possible role of the leg gap genes *homothorax, dachshund* and *Distal-less* in specifying antenniform leg fate, by characterizing their expression pattern in developing embryonic stages of *P. marginemaculatus* using newly established procedures for colorimetric whole mount in situ hybridization.

**Figure 1:**
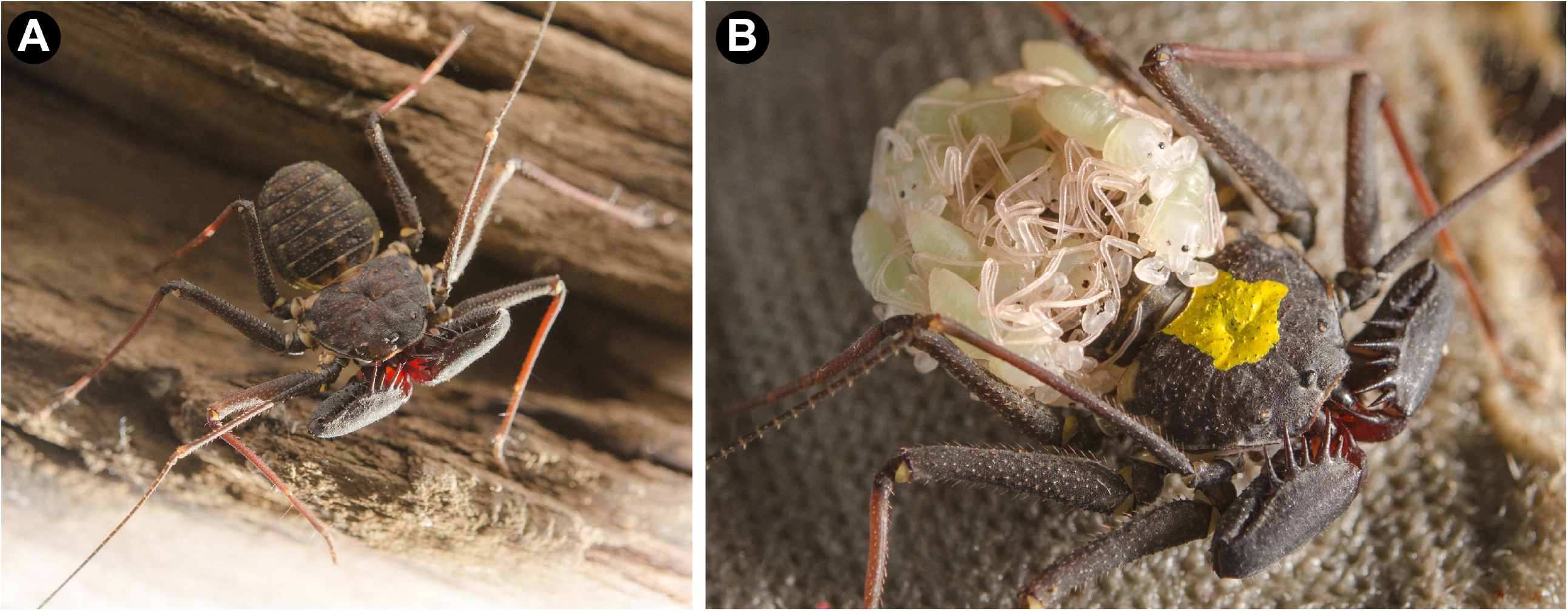
*Phrynus marginemaculatus* as a model to study Amblypygi development. A: adult male *P. marginemaculatus*, dorsal view. B: adult female P. *marginemaculatus*, carrying praenymphs on the opisthosoma. The yellow ink is used to mark the females in the colony.

## Results

### Overview of embryogenesis in Amblypygi

A characterization of the postembryonic development of *P. marginemaculatus* is available in the works of Weygoldt [71,72]. The following summary is after Weygoldt [49], complemented with observations from the present study. We note that the timing of embryonic stages is approximate, as the temperature in a previous work [49] was kept in a range of 24–27°C. Details on the culture and collection of embryos are described in *Methods*.

In our colony, a sample of 18 clutches from different females had an average of 19 eggs (min: 8 eggs; max: 39 eggs). Each egg is ovoid and measures 1.2–1.4 mm at the major axis. The yolk granules are light green, and fixation turns them light yellow. Upon secreting the brood sac and eggs into the ventral side of the opisthosoma, the brood sac is transparent, but gradually hardens and becomes dark brown in the course of two days. Around 6 dAEL, a blastoderm is formed. At 7–8 dAEL, the yolk is more condensed, a perivitelline space is formed, cells move into the yolk presumably in the region of the future blastopore. At 8–9 dAEL a blastopore is recognizable on the dorsal surface of the embryo, and a germ disc is formed. Towards 10 dAEL, the germ band starts to take form, the blastopore region is displaced posteriorly, and the first signs of segmentation of the prosoma are visible. A presumable cumulus, a group of migrating cells with dorso-ventral organizing properties, was reported by Weygoldt (1975) at this stage, but was not investigated by us. At 11-12 dAEL, the nascent prosomal segments elongate anteriorly and by 13-14 dAEL the segmented germ band occupies the ventral hemisphere of the egg. The limb buds of prosomal appendages begin to appear (Fig. 2A). The process of reversion begins, with a sagittal split of the germ band (Fig. 2A). From this point, until 19 dAEL, the ventral sulcus progressively enlarges with the dorsal displacement of both halves, and then begin to ventrally fuse again starting anteriorly (Fig. 2A–E). The head lobes on 15 dAEL have a circular outline and uniform ectoderm (Fig. 2F). From 15–19 dAEL, ectodermal cells invaginate along the ventral midline and head lobes (neural precursor cells), which can be seen as evenly spaced dark dots (Fig. 2F–H). Concomitantly, the anterior rim of the head lobes forms semilunar groves (i.e., folds over itself) (Fig. 2F–H). Opisthosomal segments 1–12 sequentially appear from the posterior growth zone (Fig. 2I–K). At 19 dAEL, the yolk moves into the caudal papilla (growth zone) and the opisthosoma flexes ventrally (Fig. 2K, N). The first pair of legs (future antenniform legs) is already longer than the other appendages in an early limb bud stage (15 dAEL), and more clearly in subsequent stages (Fig. 2L–N). A protuberance on the proximal end of the second walking leg pair (L2), the nascent lateral organ (osmoregulatory embryonic organ), is discernible at 16 dAEL and later stages, and becomes reniform (Fig. 2L–N). From 20–21 dAEL, the embryo begins to secrete a cuticle, which is darker on the fully formed lateral organ. The head lobes have completely folded onto themselves, and the distal segment of the chelicera is sclerotized. The embryo then ruptures the egg shell, and is now termed deutembryo (Fig. 2O–Q). Deutembryos remain in the brood sac for further 70 days without discernible external changes. The primordia of the eyes are formed towards the end of deutembryo development (70–106 dAEL), and eye spots of the median and lateral eyes become visible. From 90-106 dAEL the deutembryos molt into praenymphs, leave the brood sac and climb onto the adult female’s back (Fig. 1B). For a histological description of gastrulation, of the mesoderm, brain and nervous system, digestive system and gonad formation, we refer the reader to the exquisite original descriptions of Weygoldt [49].

**Figure 2:**
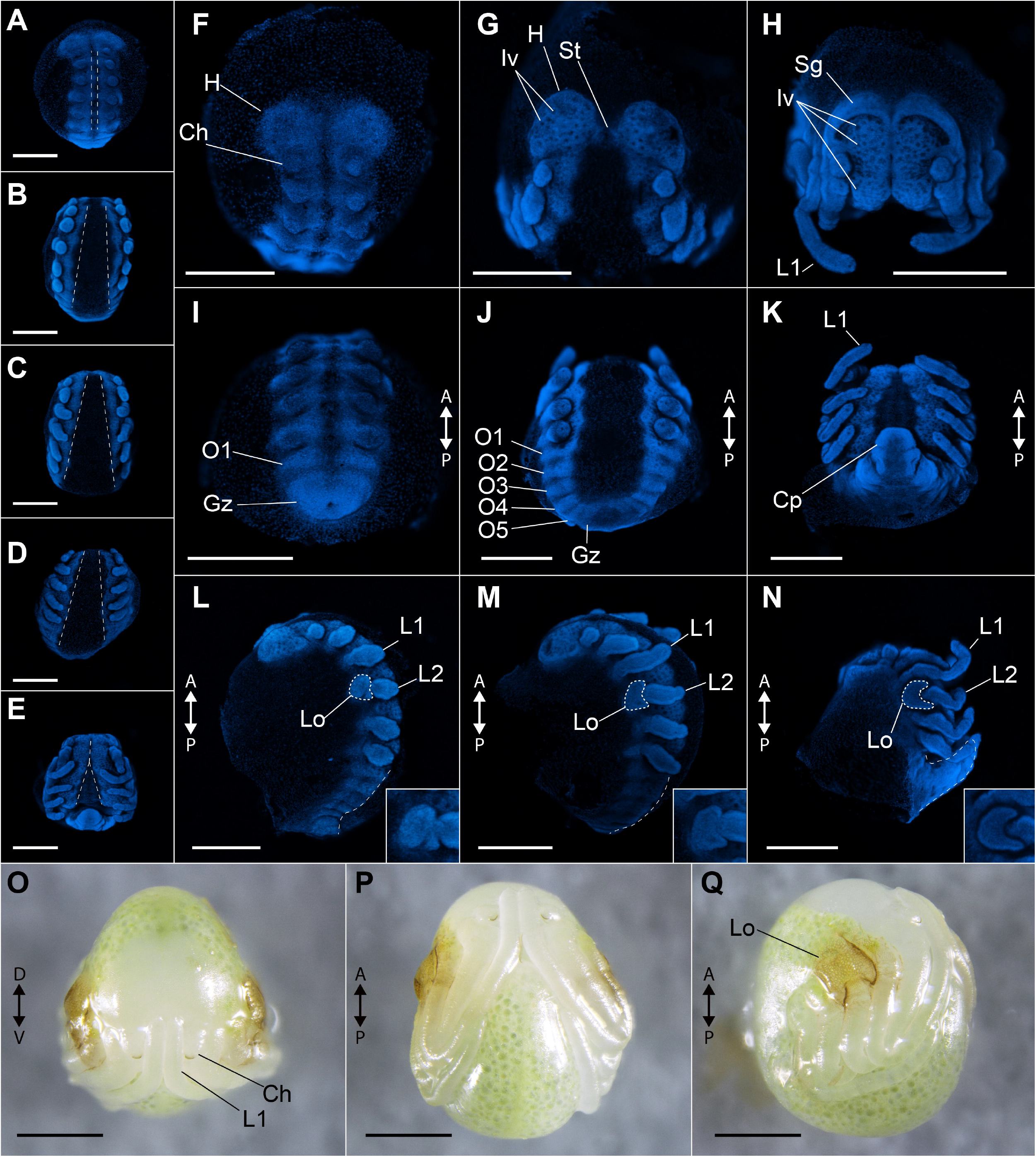
Overview of the embryonic development of *Phrynus marginemaculatus*. Z-stack automontage pictures of embryos imaged with nuclear staining (Hoechst) fluorescence. A–E: five embryonic stages, in ventral view, spanning the elongation of appendages, and showing germ band inversion and development of the ventral midline. F–H: head development, frontal view. F: undifferentiated head lobes. G: head lobe with neuron precursor cell invaginations. H: head lobes with anterior rim folding (semi-lunar grooves). I–K: opisthosomal development, posterior view. I: embryo with one opisthosomal segment. J: embryo with five opisthosomal segments. K: embryo ventral curved opisthosoma and most opisthosomal segments formed (number unclear). L–N: lateral organ formation and opisthosomal curving, lateral view. L: lateral organ as an undifferentiated bud proximal in leg II. M: lateral organ with a discreet “C” shape, and opisthosomal segments curved up. N: lateral organ in its final “kidney” shape, and opisthosoma curved ventrally. Insets in L–N show the lateral organ (Lo), which is outlined with dashed lines in the respective main panel. Dashed lines on the posterior regions of the embryos in L–N mark the curvature of the opisthosoma. O–Q: deutembryo, in frontal, ventral and lateral view respectively. Ch: chelicera; Gz: growth zone; H: head lobe; Iv: neural precursor cell invagination; St: stomodeum. Scale bars: 500 μm.

### *Phrynus marginemaculatus* embryonic transcriptome

In order to provide a comprehensive resource for Amblypygi to facilitate developmental genetic study and comparative genomics, we assembled an embryonic transcriptome spanning different embryonic stages. The assembly of *P. marginemaculatus* transcriptome resulted in 544.372 transcripts composed of 277.432.101 bp and N50 of 440 bp. BUSCO scores, which reflect the presence and number of copies of widely conserved single-copy genes (BUSCO-Arthropoda; n=1066), indicated 94.2% completeness with 5.9% of genes duplicated. From the total number of transcripts (including isoforms), we predicted 37,540 peptide sequences with the Trinotate pipeline with a top blastp hit in Swiss-Prot database. A total of 801 of these peptides (2.1%) had a top hit to non-metazoan proteins, and presumed microbial contaminants. The total number of predicted proteins is higher in comparison with the predicted protein set from the genome of the spider *Parasteatoda tepidariorum* (27,900) and the scorpion *Centruroides sculpturatus* (30,456) [6], which is attributable to the typical fragmentation of genes in such transcriptomic assemblies.

To investigate the possibility of systemic gene duplication in Amblypygi and further test the transcriptome completeness, we searched for all ten Hox genes, emphasizing discovery of paralogs known from the genomes of the spider *Parasteatoda tepidariorum* (except for *fushi tarazu*) and the scorpion *Centruroides sculpturatus* (except for *Hox3*) [6,26,32]. One of the *Hox3* copies of *P. tepidariorum* has a highly divergent homeodomain sequence, constituting an unusual long branch in our phylogenetic analysis (Additional file 1, Fig. S1). The same is true for the *Hox3* ortholog *bicoid* of *Drosophila melanogaster* (Additional file 1, Fig. S1). We therefore removed both paralogs from our main analysis. We found orthologs of all ten Hox genes in *Phrynus marginemaculatus*, namely *labial, proboscipedia, Hox3, Deformed, Sex combs reduced, fushi tarazu, Antennapedia, Ultrabithorax, abdominal A* and *Abdominal B* (Fig. 3A–B). We found evidence that all Hox genes, except *proboscipedia* are assembled in two genes, and the predicted proteins sequences overlap and have amino acid differences in *P. marginemaculatus* transcriptome (Fig. 3B). A *Pmar-Dfd* and *Pmar-abdA* were assembled in a third gene, but their non-overlapping sequences on the alignment and nearly identical amino acid sequence to the respective paralog 2 indicate that they are fragmented assemblies of the same copy. We additionally analyzed two embryonic transcriptomes of *Charinus* whip spiders recently generated by us [40]. For *C. israelensis*, we discovered orthologs of all ten Hox genes, with two genes for all cases except *ftz* and *Antp* (Fig. 3A–B). A third gene was annotated as *Cisr-Dfd*, but we interpret it as an assembly error, given its largely non-overlapping sequence with respect to the other *Cisr-Dfd* sequences. For *C. ioanniticus*, we discovered one gene each for *ftz, Antp, Ubx, AbdB*, and two genes each for *Hox3, Dfd*, and *Scr*. We did not detect orthologs of *lab, pb* and *AbdB* in *C. ioanniticus* transcriptome (Fig. 3A–B).

**Figure 3:**
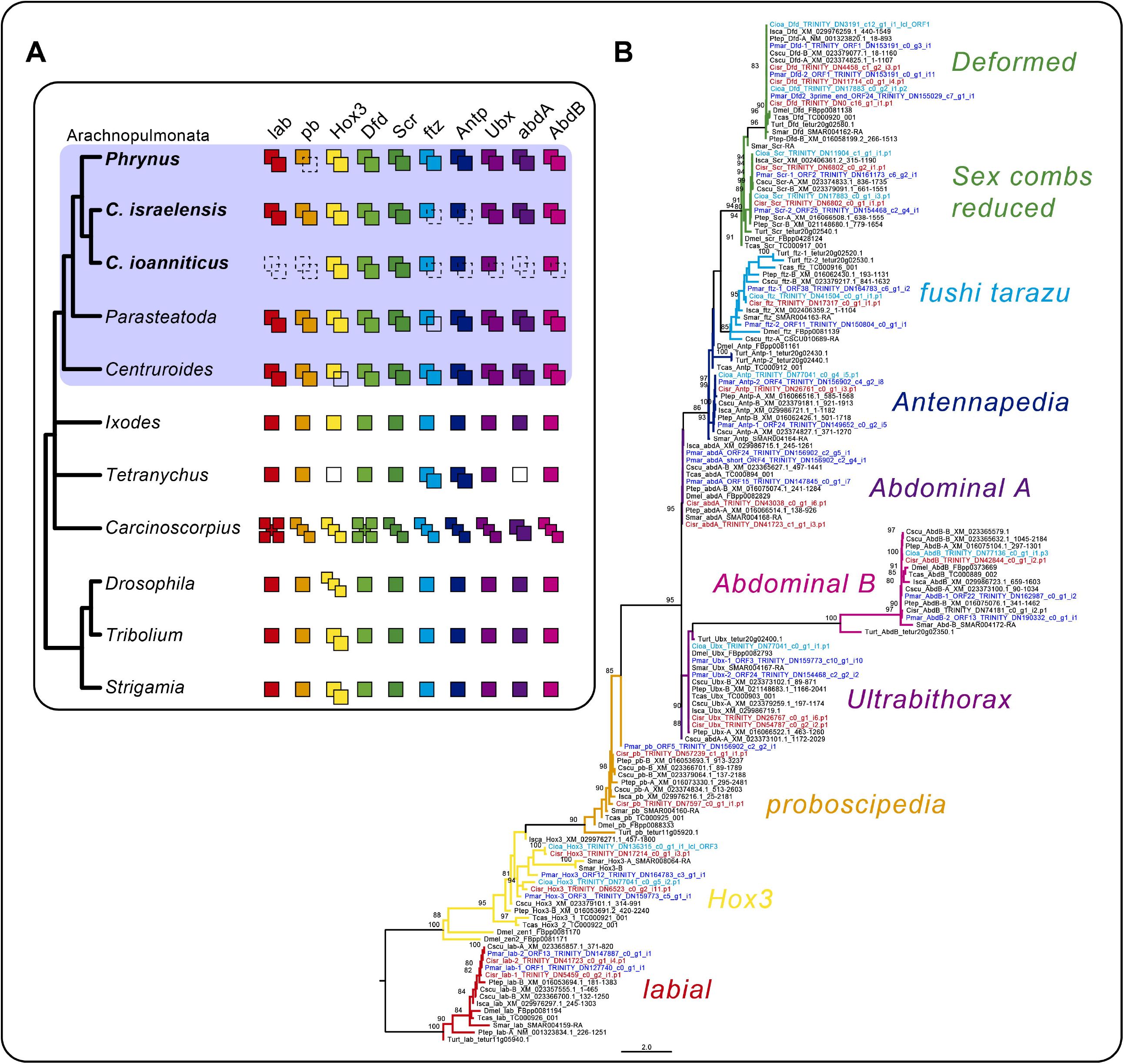
Hox gene duplications in Amblypygi. A: Schematic representation of the number of copies of each of the ten Hox genes in the selected arthropod terminals. Dotted squares in the amblypygids *Phrynus marginemaculatus, Charinus israelensis*, and *C. ioanniticus* indicate absence from the transcriptome. Empty squares in *Parasteatoda* and *Tetranychus* indicate absence from the genome. Number of Hox genes in *Carcinoscorpius rotundicauda* after [35] (but see [102] for evidence of additional copies). B: Tree topology inferred from maximum likelihood analysis of a conserved region (71 amino acid characters) using the same terminals of the schematics (ln L = −3442.810). Numbers on the notes are ultrafast bootstrap resampling frequencies (only >80 shown). Species: *Phrynus marginemaculatus* (Pmar); *Charinus israelensis* (Cisr); *C. ioanniticus* (Cioa); *Parasteatoda tepidariorum* (Ptep); *Centruroides sculpturatus* (Cscu); *Ixodes scapularis* (Isca); *Tetranychus urticae* (Ttur); *Strigamia maritima* (Smar); *Drosophila melanogaster* (Dmel); *Tribolium castaneum* (Tcas). *lab*: *labial*; *pb*: *proboscipedia*/*maxillopedia*; *Hox3*: *Hox3*/*zerknullt*/*z2*; *Dfd*: *Deformed*; *Scr*: *Sex combs reduced*; *ftz*: *fushi tarazu*; *Antp*: *Antennapedia*/*prothorax-less*; *Ubx*: *Ultrabithorax*; *abdA*: *abdominal A*; *AbdB*: *Abdominal B*.

These absences almost certainly reflect incompleteness of transcriptomic assemblies, and cannot be equated with true losses. We therefore consider the union of the Hox gene counts across the three species as the reconstruction of the whip spider ancestral Hox complement. Our results suggest that the ancestral Hox complement in Amblypygi consists of 20 genes (two copies of every Hox gene), a trait they share with spiders and scorpions. While a genome is not available for whip spiders, the phylogenetic placement of this order, as well as the pattern of Hox duplication, strongly suggests that whip spiders share the condition of two Hox clusters observed in spider and scorpion genomes.

### Gene expression analyses in *Phrynus marginemaculatus*

All three methods we trialled for fixing embryos of *P. marginemaculatus* yielded embryos which are suitable for in situ hybridization (ISH). However, fixation of intact embryos in formaldehyde/heptane phase sometimes results in the vitelline membrane adhering to the embryo. Upon dissection of the vitelline membrane for ISH, parts of the embryo are commonly lost. Piercing a hole in the embryo for fixation in formaldehyde achieves a similar result, and is therefore not recommended. Dissecting the vitelline membrane under PBS and immediately fixing the embryos in formaldehyde yielded the best results. We did not detect any difference in the signal-to-background ratio of colorimetric ISH between embryos fixed for 2 h or 24 h. To test gene expression assays and investigate the genetic patterning of antenniform legs, we selected the leg gap genes *Distal-less, dachshund* and *homothorax*, which constitute the subset of PD axis patterning genes associated with functional datasets in other arachnid model species [27,30,73].

### *Distal-less* identification and expression

Similar to other arachnopulmonates and arthropods in general, a single *Distal-less* gene, *Pmar-Dll*, was found by similarity searches and its orthology to other metazoan *Distal-less* genes was confirmed with a phylogenetic analysis (Additional file 2, Fig. S2). *Pmar-Dll* is strongly expressed in all developing appendages, from early limb bud stage (15 dAEL) and is localized to the distal portion of the chelicera, pedipalp and legs throughout leg elongation (Fig. 4A–I; Fig. 5A–L). In 17-18 dAEL embryos, the expression is heterogeneous and forms bands of stronger expression on the forming podomeres of the pedipalps and legs (Fig. 5H–L). We did not detect expression on ventral midline (*contra* [28] and[16]), or in the opisthosoma, and observed only diffuse expression on the head lobes of the stages surveyed. Expression on L1 of 16-17 dAEL embryos is subtly stronger on the proximal part of the expression domain (Fig. 5C–C’), but we did not detect clear differences in younger or older embryonic stages. The proximal and distal boundaries of strong *Distal-less* expression do not differ between L1 and the other legs, spanning two-thirds of the legs from the tip of the tarsus and excluding the most proximal third (Fig. 5C, 5I).

**Figure 4:**
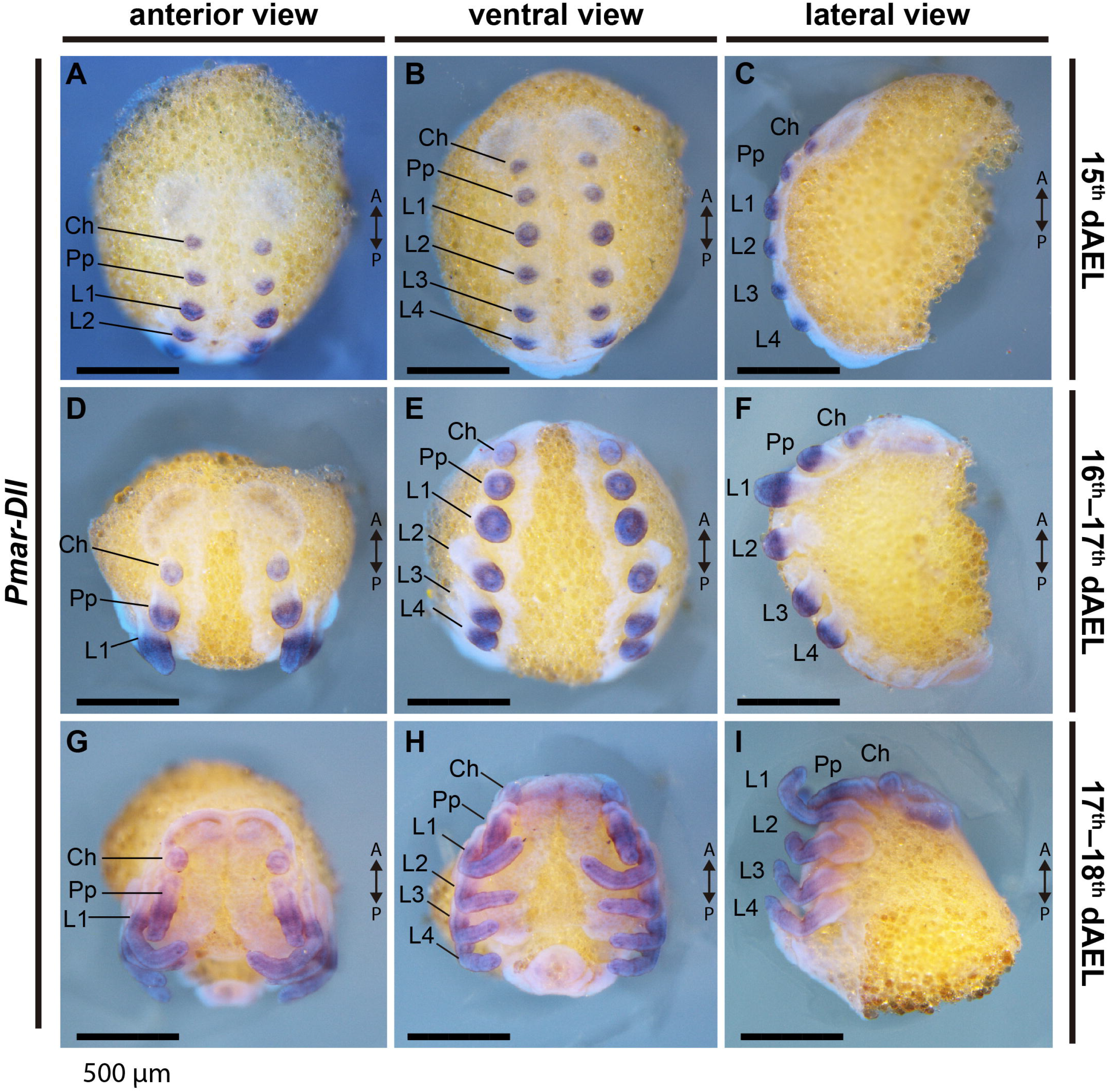
*Pmar-Distal-less* (*Pmar-Dll*) colorimetric in situ hybridization. Whole mount bright field images (Z-stack automontage) overlaid with nuclear staining (Hoechst). Ch: chelicera; Pp: pedipalp; L1–4: leg 1–4; A/P: anterior/posterior; dAEL: days after egg laying. Scale bars: 500 μm.

**Figure 5:**
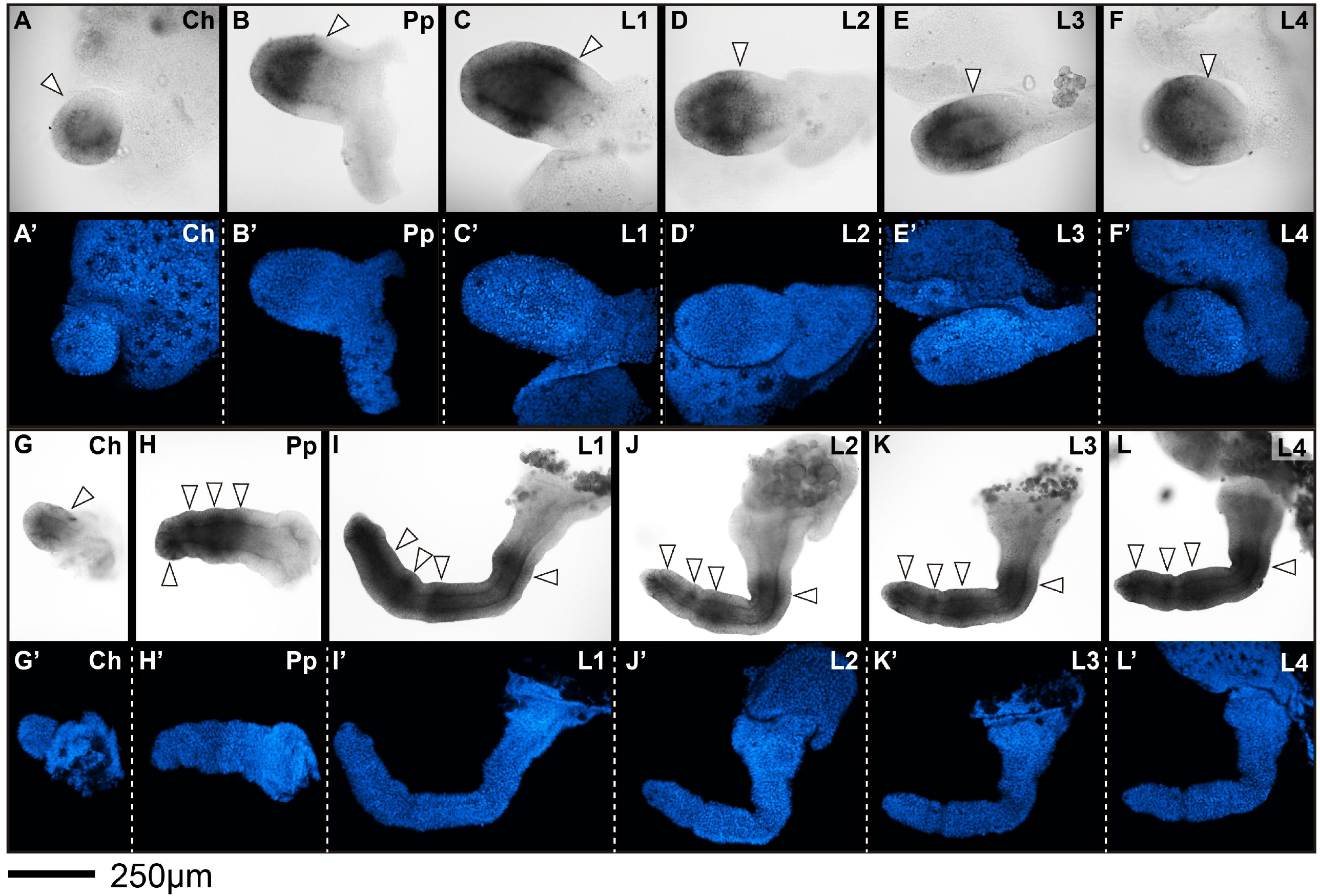
*Pmar-Distal-less* (*Pmar-Dll*) colorimetric in situ hybridization, appendage dissections in lateral view. Distal is to the left. A–F, G–L: bright field. A’–F’, G’-L’: nuclear staining (Hoechst), confocal Z-stack maximum projection. A–F: 16–17 dAEL embryo. G–L: 17-18 dAEL embryo. J and J’ have been vertically reflected to adjust orientation. Distal is to the left. Arrowheads in A–F indicate proximal boundary of expression. Arrowheads in G–L indicate stronger domains of expression. Ch: chelicera; Pp: pedipalp; L1–4: leg 1–4. Scale bar: 250μm.

In contrast to other arachnid embryos (mygalomorph spiders [69]; araneomorph spiders [67,73]; Opiliones, Sharma et al. 2012; and mites [20,74]), *Dll* expression was not detected as a strong domain in the endite of pedipalp, which is very inconspicuous at these embryonic stages. We were not able to assess *Dll* expression in the pedipalpal endite in deutembryo stages due to deposition of cuticle.

### *dachshund* identification and expression

*dachshund* is typically present as a single copy in Arthropoda, but is found independently duplicated in Arachnopulmonata and Xiphosura [38,75]. We found two genes with high similarity to *Dmel-dac* and confirmed their identity as dachshund paralogs with a phylogenetic analysis (Additional file 3, Fig. S3). *Pmar-dac1* and *Pmar-dac2* are each nested in a clade with paralogs 1 and 2 of other arachnopulmonates, which appears as independent from horseshoe crab duplications (Additional file 3, Fig. S3).

*Pmar-dac1* is strongly expressed as broad median band in the pedipalp and the four pairs of legs (Fig. 6A–F). This sharp mid band excludes the most proximal and the distal domains of these appendages (Fig. 7). No staining was detected on the chelicera (Fig. 6A–F; 7A, E). Additionally, expression occurs in three domains in the head lobes and strongly on the ventral midline of the prosoma. The continuous expression on the ventral midline stops before the opisthosoma (Fig. 6A–F).

**Figure 6:**
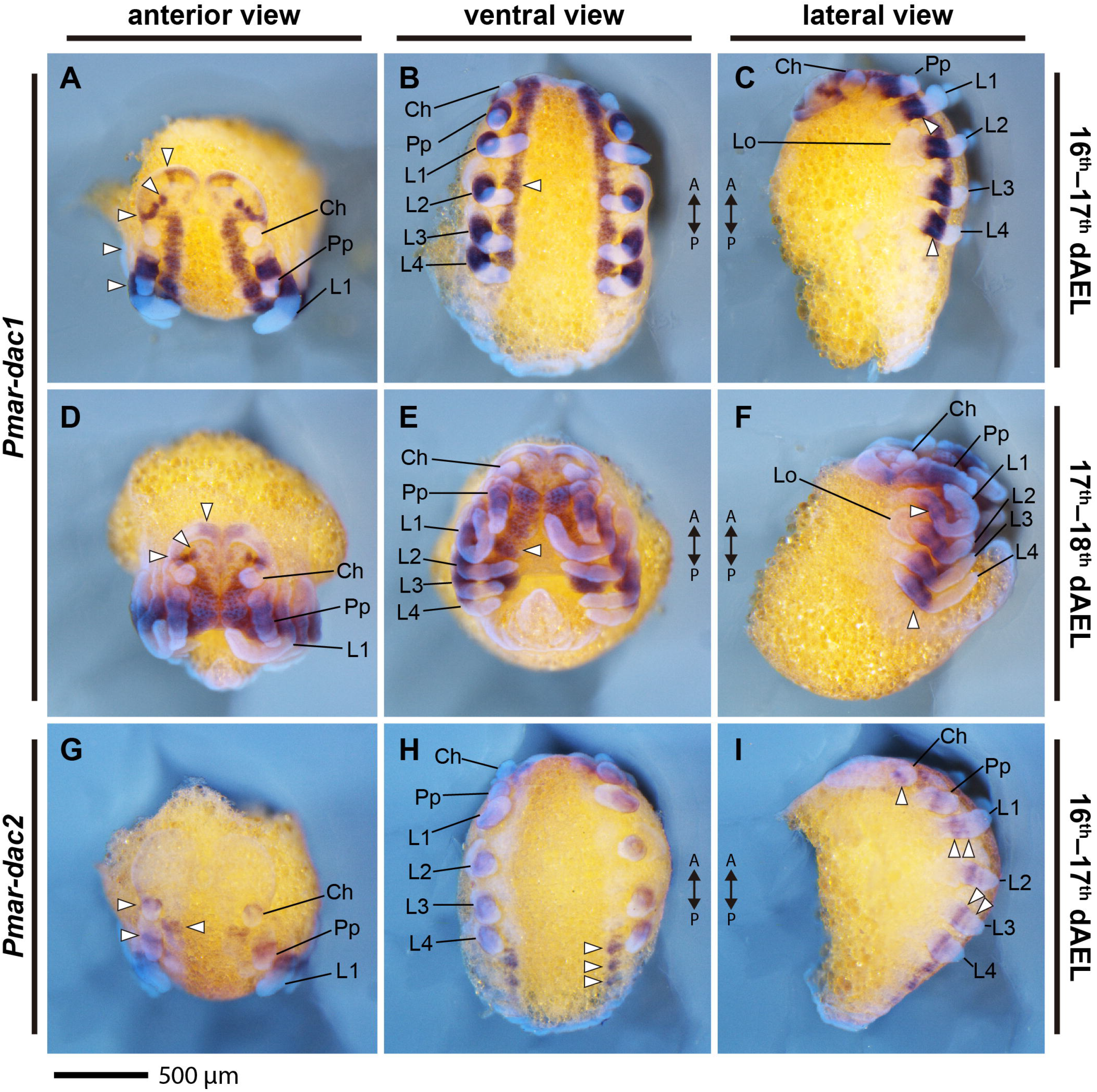
*Pmar-dachshund 1* (*Pmar-dac1*) (A–F) and *Pmar-dachshund 2* (*Pmar-dac2*) G–I) colorimetric in situ hybridization. Whole mount bright field images (Z-stack automontage) overlaid with nuclear staining (Hoechst). Ch: chelicera; Pp: pedipalp; L1–4: leg 1–4; Lo: lateral organ; A/P: anterior/posterior; arrowhead: expression domain; dAEL: days after egg laying. Scale bars: 500 μm.

**Figure 7:**
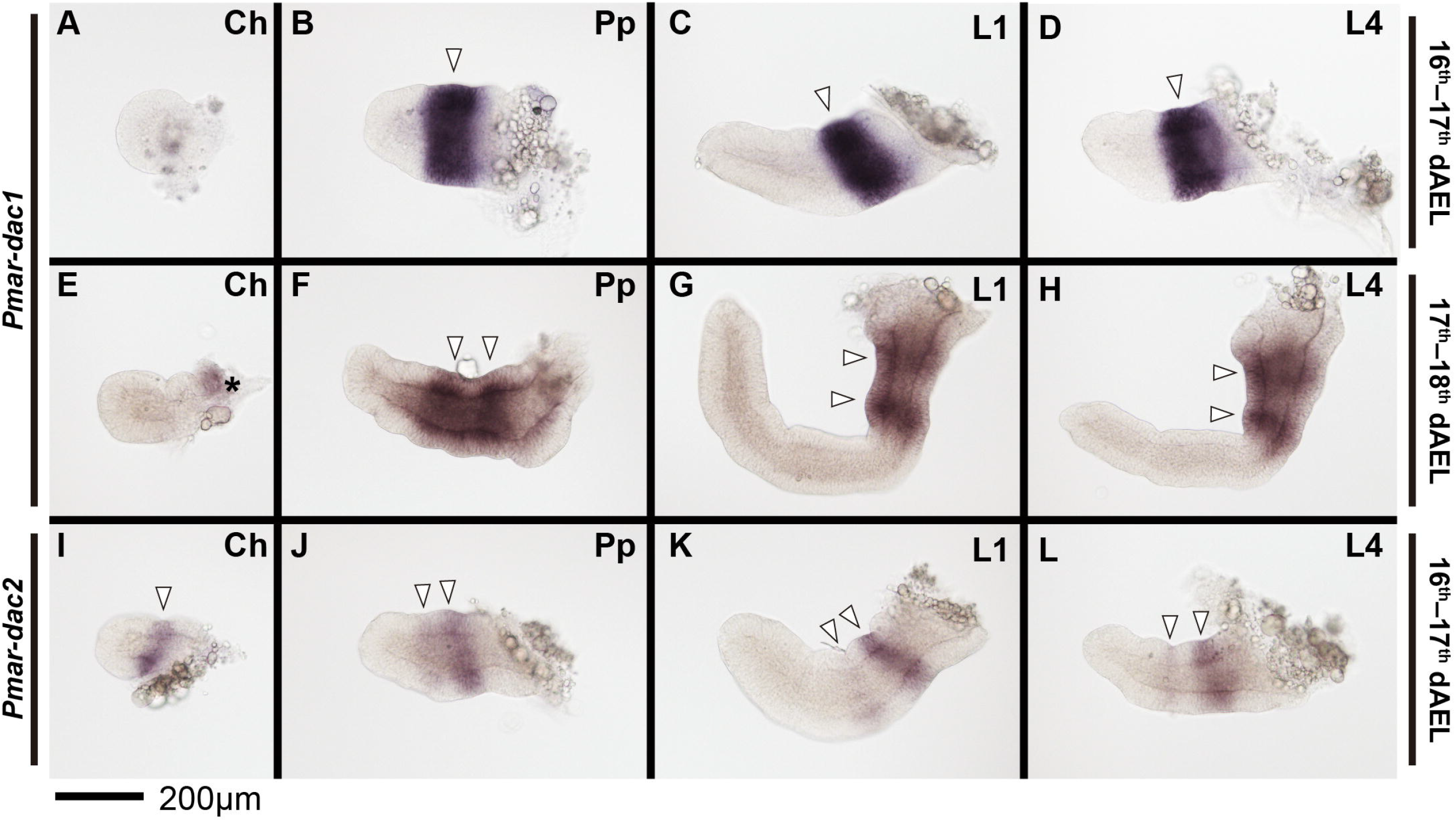
*Pmar-dachshund 1* (*Pmar-dac1*) (A–H) and *Pmar-dachshund 2* (*Pmar-dac2*) (I–L) colorimetric in situ hybridization, appendage dissections in lateral view. Distal is to the left. White arrowhead indicates discreet domains of expression. Asterisk in “E” marks expression from the lateral margin of the headlobe. Ch: chelicera; Pp: pedipalp; L1, 4: leg 1, 4. dAEL: days after egg laying. Scale bar: 200 μm.

*Pmar-dac2* is expressed as ring at the proximal region of chelicera, pedipalp and legs (Fig. 6G–I). On 16-17 dAEL embryos, a second thinner ring of expression is also present on the pedipalps and legs (Fig. 6G–I; 7I–L). Expression domains also occur on the neuromere of L1 and pedipalp, being larger on the pedipalpal neuromere (Fig. 6G). Three spot domains also occur on O1, O2 and O3 segments (Fig. 6H–I).

### *homothorax* identification and expression

Similar to the case with arachnid *dachshund, homothorax* is also found duplicated in arachnopulmonates [32,39]. We annotated two genes in *P. marginemaculatus* transcriptome and the phylogenetic analysis shows that each copy is nested in two clades with the arachnopulmonate paralogs (Additional file 4, Fig. S4).

*Pmar-hth1* is conspicuously expressed on the developing appendages throughout the stages studied (15-18 dAEL; Fig. 8–9). On 16-17 dAEL embryos, *Pmar-hth1* is expressed more strongly on the proximal parts of the chelicera, pedipalp, and legs, and is absent in a small distal domain (Fig. 9A–D). On 17-18 dAEL, expression in all elongated appendages is similar, and spans most of the appendage length except for a small distal territory (Fig. 9E-H). Additionally, three stronger stripe domains occur distal to the coxa of pedipalp and legs. Expression in L1 and other legs is similar in the studied stages, save for a slightly expanded distal domain on the proximal tarsus of L1 (Fig. 9G–H). Expression on the ventral ectoderm and head lobes is weak and homogenous in early stages, and becomes stronger in 17-18 dAEL (not shown)

**Figure 8:**
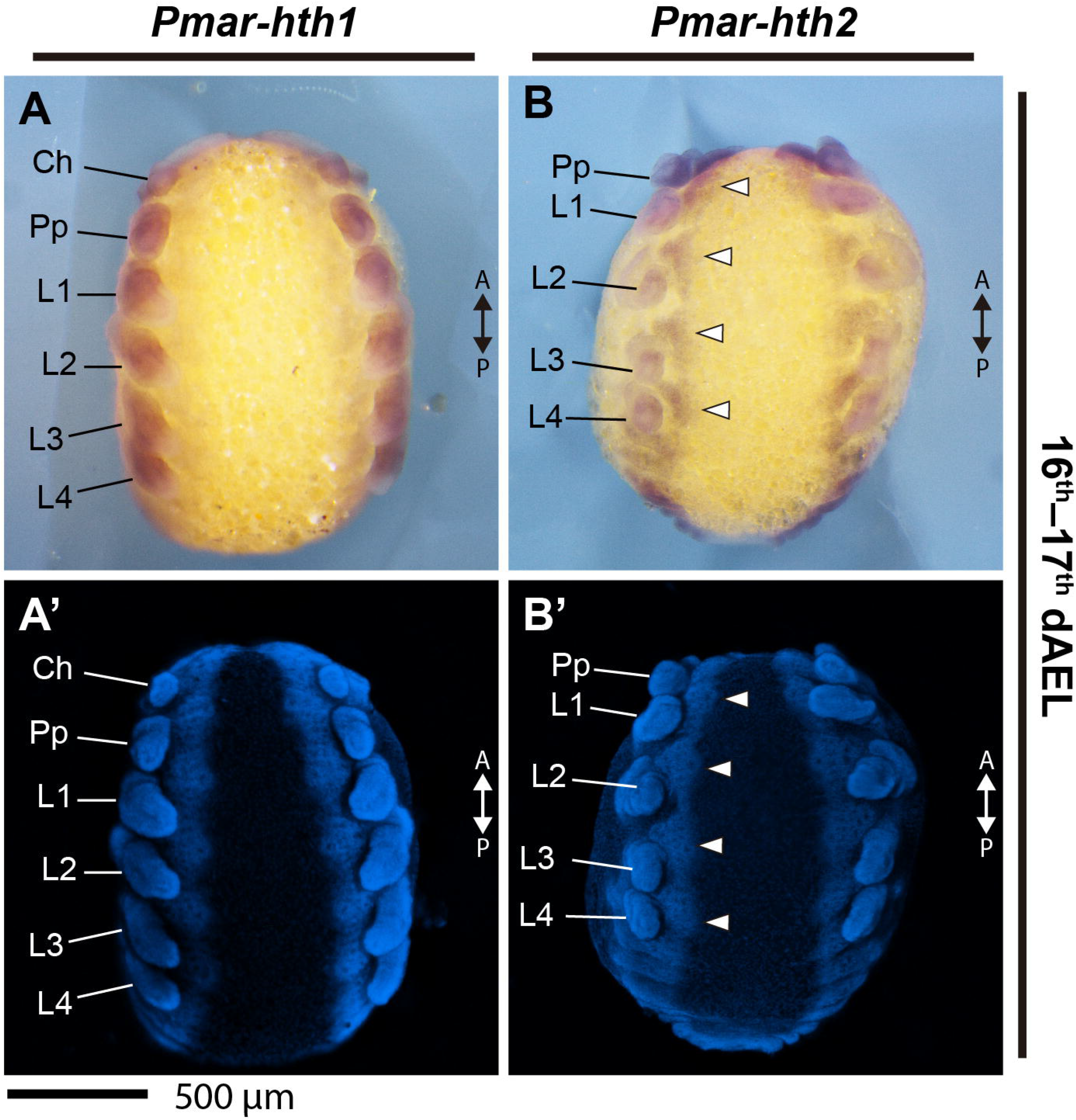
*Pmar-homothorax 1* (*Pmar-hth1*) (A–A’) and *Pmar-homothorax 2* (*Pmar-hth2*) (B– B’) colorimetric in situ hybridization, ventral view. A, B: whole mount bright field images (Z-stack automontage). A’, B’: nuclear staining (Hoechst) (Z-stack automontage). Ch: chelicera; Pp: pedipalp; L1–4: leg 1–4. A/P: anterior/posterior; arrowhead: expression domain; dAEL: days after egg laying. Scale bar: 500 μm.

**Figure 9:**
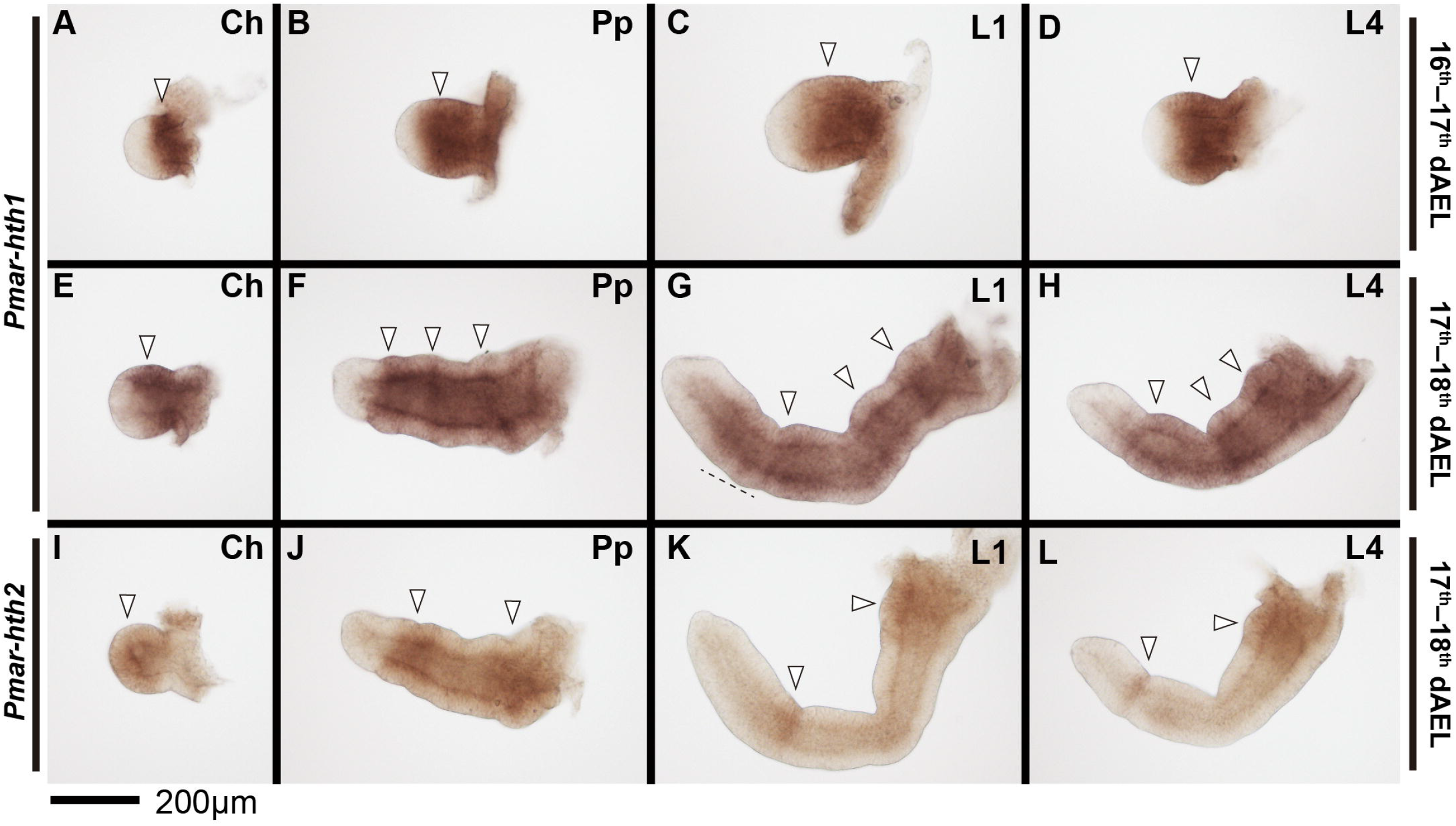
*Pmar-homothorax 1* (*Pmar-hth1*) (A–H) and *Pmar-homothorax 2* (*Pmar-hth2*) (I– L) colorimetric in situ hybridization, appendage dissections in lateral view. Distal is to the left. White arrowhead indicates discreet domains of expression. Dotted line in “G” marks expanded distal expression domain. Ch: chelicera; Pp: pedipalp; L1, 4: leg 1, 4. dAEL: days after egg laying. Scale bar: 200 μm.

*Pmar-hth2* is expressed in the developing appendages, but is reduced in intensity and different in pattern in comparison to *Pmar-hth1* (Fig. 8B–B’). *Pmar-hth2* spans the whole chelicera (Fig. 9I). Expression on the pedipalp spans most of the appendage, excluding only a distal territory (Fig. 9J). In addition, there is a proximal and a sub terminal strong domain (Fig. 9J). Expression on L1 and other legs is identical in all stages studies. *Pmar-hth2* expression on the legs is consistently weaker in comparison to the pedipalp, but likewise possess a proximal and a distal sub-terminal domain (Fig. 9K–L). Expression on the ventral ectoderm, opisthosoma and head lobes is ubiquitous (Fig. 8B).

## Discussion

### Genomic resources for whip spiders corroborate widespread retention of duplicates across Arachnopulmonata

Changes in Hox gene expression, regulation and targets are linked to changes in morphology and Hox gene duplications have also been implicated in morphological innovations [76-79]. Duplication in the Hox cluster is common in the vertebrates, which may have up to eight Hox clusters (teleost fishes) due to successive rounds of genome duplications [76]. On the other hand, genome and Hox cluster duplications are considered rare in Arthropoda [80]. Complete Hox cluster duplication in the phylum has only been reported in horseshoe crabs (Xiphosura) [34,35,81,82], and in spiders and scorpions (presumed ancestral in Arachnopulmonata). The phylogenetic placement of Amblypygi makes this group an opportune system to test the downstream prediction that *P. marginemaculatus* retains and expresses Hox paralogs shared by spiders and scorpions. In accordance with this hypothesis, we discovered that the whip spider *P. marginemaculatus* presents homologs of all ten Hox genes, and all excepting *proboscipedia* appear to be duplicated, as supported by our alignment and phylogenetic analysis. Our survey in the two *Charinus* transcriptomes confirmed the duplication of several Hox genes, and additionally revealed that two copies of *proboscipedia* (absent in *P. marginemaculatus*) are present at least in one species. Our results suggest that Amblypygi retain a complete duplicated Hox complement with 20 genes. Similarly, a recent transcriptomic survey in *Charinus acosta* and *Euphrynichus bacillifer* reported duplications of nearly all Hox genes (excepting *proboscipedia*), as well as Wnt and Frizzled genes [83]. Taken together with the recent discovery of duplicated RDGN genes in *Charinus*, our results suggest that Amblypygi retain ohnologs of numerous developmental patterning genes, as a function of an ancient, shared whole genome duplication.

A second line of evidence of a shared genome duplication event comes from conserved expression patterns of leg gap gene paralogs. It has recently been shown that duplicates of leg patterning genes *extradenticle, optomotor blind, dachshund* (*dac*) and *homothorax* (*hth*) present one copy with an expression pattern resembling a stereotypical panarthropod domain, and a second copy with a divergent expression, with these two sets of expression patterns shared by a spider and a scorpion [75]. For instance, *dac1* of surveyed arachnopulmonates retains the plesiomorphic expression pattern of a broad median band in the appendages, the condition observed in the single-copy *dac* of all non-arachnopulmonate arachnids surveyed to date [20,28], whereas *dac2* is expressed as a proximal and a medial domain at the patella-tibia boundary [38,67,69,75]. Similarly, *hth1* retains a plesiomorphic expression pattern on the proximal-median appendage excluding the tarsus, whereas *hth2* presents a strong short proximal domain and stripes of expression at podomere boundaries [39,67,69,75]. In the whip spider *P. marginemaculatus*, we indeed recovered two copies of *dachshund* and *homothorax*, with paralogs nested in two clades of arachnopulmonate copies with high nodal support (Additional file 3, Fig. S3; Additional file 4, Fig. S4). Moreover, *Pmar-dac* and *Pmar-hth* have paralog-specific expression patterns shared with spiders and a scorpion: *Pmar-dac1/2* and *Pmar-hth1/2* conserve the plesiomorphic expression pattern of arachnids, while copy 2 retains the arachnopulmonate-specific pattern. These data, together with the evidence from Hox gene duplications and the topologies of the *dac* and *hth* gene trees, provide compelling support for genome duplication as a synapomorphy of Arachnopulmonata.

### A *dachshund2* domain in the two-segmented clasp-knife chelicera of *P. marginemaculatus*

Expression of *Pmar-Dll, Pmar-dac1/2* and *Pmar-hth1/2* on the appendages is largely similar to spiders and scorpions (see above) [38,39,67-69,75]. A notable difference in the chelicera of the whip spider is the retention of proximal expression of one of the *dachshund* paralogs. In arachnids with three segmented chelicerae, such as the daddy-long-legs *Phalangium opilio, dac* is expressed in the proximal segment of the appendage [28]. The same is true for the three-segmented chelicera of scorpions, in which both *dac1* and *dac2* are expressed in the proximal segment [75]. In the two-segmented chelicera of spiders, *dac1* is not expressed in the chelicera save for a non-ectodermal faint staining in late embryonic stages [38,67-69,84]. *dac2* of spiders is also largely not expressed in the chelicera, save for a small ventral domain at the base of the chelicera of *Parasteatoda tepidariorum* and *Pholcus phalangioides* [38]. It has been proposed that the loss of a proximal *dac* domain is associated with the loss of the proximal segment in the chelicera of spider and other arachnids. This hypothesis is supported by functional data in the daddy-long-leg *P. opilio*: knockdown of the single-copy *dac* resulted in the loss of the proximal segment of the chelicera (as well as mid-segment truncations in pedipalps and legs) [27]. Similar to spiders, the two-segmented chelicera of the whip spider *P. marginemaculatus* does not express *Pmar-dac1*. Nonetheless, a clear complete ring of proximal expression was detected for *Pmar-dac2*.

In contrast to the classic leg gap phenotype incurred by knockdown of the ancestral *dac* copy in the harvestman, knockdown of *dac2* in the spider *P. tepidari*orum resulted in the loss of the patella-tibia segment boundary in the pedipalp and walking legs; effects of this knockdown on the chelicera were not reported by the authors, nor is the function of *dac1* known in *P. tepidariorum* [38]. Given the interpretation that *dac2* has a derived role in patterning a segment boundary in the spider, the incidence of the *Pmar-dac2* stripe in the middle of the whip spider chelicera (and the absence of *Pmar-dac1* in the chelicera) is tentatively hypothesized here to constitute evidence of a segment boundary patterning activity that separates the basal and distal cheliceral segment of Amblypygi (rather than evidence of a proximal segment identity in whip spiders that is homologous to the proximal cheliceral segment of groups like harvestmen and scorpions). Exploring this hypothesis in future will require the establishment of RNAinterference tools in *P. marginemaculatus*.

### Antenniform leg distal allometry is specified early in development

To investigate a possible role of changes in leg gap genes expression in the patterning of the antenniform leg pair (L1) of whip spiders, we specifically searched for changes in expression domain between embryonic antenniform leg (L1) and the posterior legs (L2–4). The expression patterns of *Pmar-Dll, Pmar-dac1/2* and *Pmar-hth1/2* are mostly identical between the antenniform leg L1 and the posterior legs. We found only a slightly expanded *Pmar-hth1* domain in the distal leg (compare Fig. 9G–H). It is unclear if this difference bears any functional significance in the differential fate of L1. Similar distal expression dynamics appear to distinguish the identity of the pedipalp and walking legs of the harvestman, as knockdown of the single copy of *hth* in *P. opilio* results in homeotic transformation of pedipalps to legs [30].

Besides the difference in sensory equipment, the antenniform leg pair of adults differ from the posterior walking legs in being much longer, and presenting numerous articles (pseudo-segments) on the tibia, metatarsus and especially the tarsus, whereas the walking legs have pseudo-segments only in the tarsus [41,53,55]. Antenniform legs may be more than 2.5 times longer than the walking legs, and this allometry in adults is mostly accounted for by the length added by the distal leg segments, which may attain more than 100 distal articles (tibiomeres + tarsomeres) [53]. Understanding the timing of the ontogenetic divergence of L1 relative to the walking legs is crucial for circumscribing target stages for investigation of the genetic patterning of this phenomenon.

Current evidence indicates that this allometry on the antenniform legs takes place both embryonically and postembryonically. The praenymph, the stage that molts from the deutembryo and climbs to the mother’s back, has the full number of tibiomeres, but not of tarsomeres. The final number of tarsomeres is added in the next molt to the protonymph [53,85], and from this moment onwards the subsequent growth is isometric [53,85]. In embryos of *P. marginemaculatus*, Weygoldt [49] observed that the primordia of L1 are the first to appear in the germ band and are already longer than the remaining prosomal appendages, an observation paralleling the allometric growth of the L2 limb bud of the harvestman *P. opilio* (wherein L2 is the longest leg and also bears the most tarsomeres) [29]. It is presently unclear at what point in development the distal podomeres of antenniform legs attain their allometry with respect to the proximal podomeres in the whip spider. In other words, are embryonic legs simply growing longer overall with respect to walking legs, or are they longer due to distal growth? The expression pattern *dachshund* in *P. marginemaculatus* early embryos reveals a further interesting aspect of this allometry. The expression of *dac* (including duplicates) in the legs of arachnids surveyed to date occurs on podomeres proximal to the tibia, so that the tibia-metatarsus-tarsus territory may be identified as the a *dac*-free domain [20,27,38,67,69,75,84]. The distal boundary of *Pmar-dac1* and *Pmar-dac2* in stages long before podomere formation reveals that the distal domain (*Pmar-dac1*-free cells) of L1 is already longer with respect to the proximal leg and the other legs (compare Fig. 7C, D). These data support the inference that the distal allometry, and by extension the antenniform leg fate, are already specified from the onset of limb bud development.

## Conclusion

The study of the embryology of Amblypygi has faced 45 years of hiatus since the seminal work describing the development of *Phrynus marginemaculatus* [49]. Here, we take the first steps in establishing *P. marginemaculatus* as a model for modern chelicerate evolutionary developmental study and for comparative genomics. We provide evidence for a shared genome duplication in Amblypygi; show that the distal allometry of antenniform legs is specified early in appendage embryogenesis; and observe that expression patterns of the leg gap genes *Distal-less, dachshund* and *homothorax* do not substantian their role in fate specification of the sensory legs of whip spiders. Future efforts to uncover the genetic underpinnings of this fascinating convergence with the mandibulate antenna should prioritize discovery of leg fate specification mechanisms in early whip spider development. Despite the establishment of a few promising arachnid models for studying chelicerate embryology, a pervasive challenge has been the establishment of toolkits for testing gene function [3,22,27,73]. Therefore, future efforts to study whip spider development must emphasize the development of tools for functional experiments in *P. marginemaculatus*.

## Methods

### Establishment of amblypygid colony in the laboratory

Adult individuals of *Phrynus marginemaculatus* were purchased from a commercial supplier collects animals in the Florida Keys (USA). The maintenance of a colony of *Phrynus marginemaculatus* (Fig. 1A–B) in the laboratory followed guidelines provided is the original developmental works of Peter Weygoldt [49,71,72] and notes on behavioral studies on this species [86-88]. While most species of Amblypygi have been reported as aggressive and better kept in separate containers [41], individuals of *P. marginemaculatus* can be kept in a stable laboratory colony in large community containers [86]. A maximum of 20 animals may be allowed to interact and mate freely in plastic boxes (67.3 × 41.9 × 32.4 cm) with a 2 cm layer of damp coconut fiber saturated with distilled water. The substrate is continuously dampened with approximately 500 mL of dH_2_O on the substrate every week. All animals are kept in an acclimatized room at 26-28°C in a controlled reversed 16L:8D light cycle to simulate summer months in Florida. To prevent flies from reproducing in this humid environment, the lid of the container is provided with a 10 × 20 cm fine mesh cloth, which also allows for air circulation. Each container is furnished with pieces of tree bark, egg carton, and retreats made from polystyrene foam or florist’s foam, to provide enough space for animals to hide and suspend themselves during molting and egg-laying. After about two weeks of setting up a terrarium, the substrate hosted a stable colony of collembolans (possibly brought by the tree bark). The co-occurrence of collembolans does not appear to be disadvantageous in any way. Collembolans are invariably observed feasting on carcasses and likely contribute to preventing mite infestations (see also [41]). Animals are fed at least twice a week with nymphs of *Acheta domestica ad libitum*. Carcasses are removed as soon as detected to prevent the spread of mites. Freshly hatched praenymphs are kept with the female, and aggregate on the back of the brooding female (Fig. 1B). The next instar, the protonymph, disperse from the female upon molting. Protonymphs are transferred to community terraria as described above. Protonymphs and subsequent instars are fed pinheads of *A. domestica* and *Gryllus bimaculatus ad libitum* three times per week until adulthood. The first adult individuals from our laboratory-bred animals were detected 8 months after hatching, which accords to what has been described by Weygoldt [49].

Terraria with adults are inspected daily for freshly molten individuals or ovigerous females, which are immediately separated from the main colony to avoid aggressive interactions. Ovigerous females carry the eggs outside the body in a brood sac attached to the ventral side of the opisthosoma [41]. Eggs are allowed to develop with the female until the desired stage for developmental work. Approximate stages may be inferred from days after egg laying (dAEL) at 26°C following the staging in [49]. For obtaining the eggs, females are anesthetized with CO_2_ and immobilized in customized plastic petri dishes (35 × 11 mm) with a central whole and foam (Fig. 10A–B). With a blunt forceps pair, the pleura of the opisthosoma embracing the eggs may be gently displaced from the brood sac, and the brood sac removed as whole (Fig. 10C). The female may be returned to the colony unharmed. The brood sac easily dissolves when immersed in a phosphate buffer solution (PBS), and eggs may be cleaned with forceps under PBS. We obtained high mortality in keeping isolated eggs developing in petri dishes with humid (wet filter paper) or dry in 26°C incubators, as mold frequently develops. However, we found out that embryos will survive and develop immersed in PBS supplemented with kanamycin (1:5000) at 26°C incubator and even hatch into deutembryos (∼20 dAEL).

**Figure 10:**
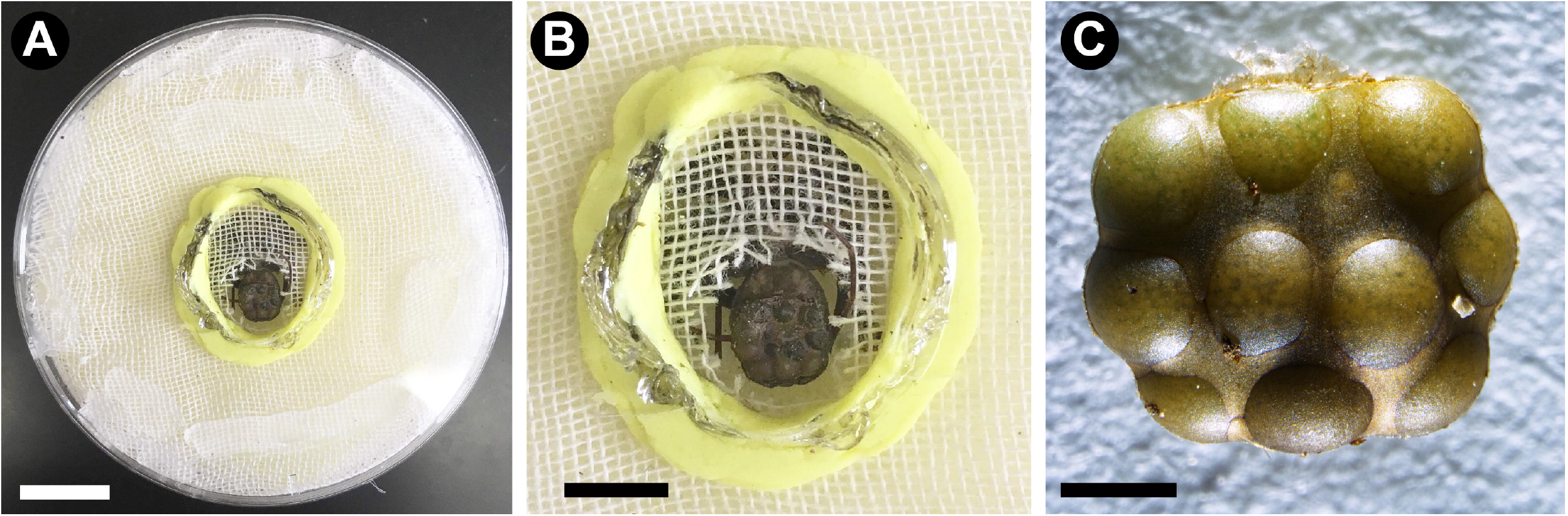
Tool for removing the brood sac from whip spiders. A: dissecting dish made of plastic petri dish and foam. B: detail of an immobilized female ready to have the brood sac dissected. C: intact brood sac dissected from the female. Scale bars: A: 1 cm (approximate); B: 5 mm (approximate); C: 1 mm.

### Phrynus marginemaculatus transcriptome assembly

We collected embryos of different embryonic stages as described above, and conducted total RNA extractions of each stage separately: hatched deutembryo (female #5, n=3; undetermined age, before eyespots), unhatched deutembryo (female #2 n=3; approx. 21 dAEL), and leg elongation stage (female #3, n=2; approx. 16-17 dAEL), late limb bud stage (female 6, n=1; 16 dAEL) and early limb bud stage (female 6, n=1; 14 dAEL) (voucher images are available upon request). Embryos were homogenized in Trizol TRIreagent (Invitrogen) and stored at –80°C until extraction. The combined total RNA of the five extractions was submitted to cDNA library preparation and sequencing in an Illumina High-Seq platform using paired-end short read sequencing strategy (2 × 100 bp) at the Center for Gene Expression, Biotechnology Center of University of Wisconsin-Madison (USA). The quality of the raw reads was assessed with FastQC (Babraham Bioinformatics). De novo assembly of compiled paired-end reads was conducted with the software Trinity v. 2.6.6 [89], using Trimmomatic v. 0.36 [90] to remove low quality reads and adaptors. Completeness of the transcriptome was assessed with the Trinity script ‘*TrinityStats*.*pl*’, and BUSCO v. 3 [91] on the longest isoforms using the database ‘Arthropoda’ and analysis of Hox genes. Whole transcriptome annotation was conducted following the Trinotate pipeline, by predicting proteins with TransDecoder v. 5.2.0 [92] and searching best hits with blastx and blastp against Swiss-Prot and Pfam databases [93].

### Gene identification

Hox genes (*labial, proboscipedia, Hox3, Deformed, Sex combs reduced, fushi tarazu, Antennapedia, Ultrabithorax, abdominal A* and *Abdominal B*), *Distal-less, dachshund*, and *homothorax* orthologs were identified from the embryonic transcriptome of *Phrynus marginemaculatus* using BLAST searches (tblastn) with peptide queries of the homologs of *Drosophila melanogaster* and *Parasteatoda tepidariorum*. The best hits were reciprocally blasted to NCBI database and further assessed with a phylogenetic analysis.

For the phylogenetic analysis of Hox genes, we selected arthropod terminals with available genomes and well annotated references: the fruit fly *Drosophila melanogaster* (FlyBase), the beetle *Tribolium castaneum* (GCA_000002335.3) (Insecta), the centipede *Strigamia maritima* (GCA_000239455.1) (Myriapoda), the tick *Ixodes scapularis* (GCA_002892825.2), the mite *Tetranychus urticae* (GCA_000239435.1), the spider *Parastetoda tepidarioum* (GCA_000365465.2) and the scorpion *Centruroides sculpturatus* (GCA_000671375.2) (Chelicerata) [6,94-97] (Additional file 5, Table S1). For the analysis of *Distal-less, dachshund*, and *homothorax* we selected terminals used in Nolan et al. [75] and included the peptide sequence for *P. marginemaculatus* candidates (Additional file 5, Table S1). Peptides were predicted with ORF Finder (NCBI), TransDecoder v. 5.5.0 [92], or obtained as CDS when possible from publicly available NCBI database. Peptide sequences were aligned using Clustal Omega [98] in the software SeaView v. 4 [99]. Gene trees were inferred under maximum likelihood using IQTREE [100] with automatic model selection (-m TEST) and 1000 ultrafast bootstrap resampling (-bb 1000). Alignments for the Hox gene analysis are available in Additional files 6–7. Genes referred throughout text refer to the longest isoform assembled by Trinity in a contig, and annotated by the procedure detailed above. Paralogs were discerned from gene fragments using as criteria the presence of largely overlapping sequences with amino acid differences.

### Fixation and in situ hybridization

Collecting freshly laid eggs and keeping them in controlled temperature with the female or incubator (26°C) allows to select desirable stages for fixation. We fixed embryos for in situ hybridization with three methods: (1) intact eggs in a phase of 4% formaldehyde/PBS and heptane overnight, after an established spider fixation protocol [3]; (2) piercing a hole in the vitelline membrane, and immersing eggs in 4% formaldehyde/PBS overnight; (3) dissecting the embryo out of the vitelline membrane and fixing in 4% formaldehyde/PBS for periods varying from 2–24h. All three procedures were conducted in a shaking platform at room temperature (∼22 °C). Following the fixation period, embryos were rinsed several times in 1x PBS/0.1% Tween-20 (Sigma Aldrich) (PBS-T), and gradually dehydrated into pure methanol. Embryos were then stored in 20 °C freezer until next procedures.

cDNA was synthetized with SuperScriptIII kit (Thermo Fisher) following the manufacturer’s instructions, using the same total RNA used to generate the embryonic transcriptome. Riboprobe templates for the genes *Pmar-dac1, Pmar-dac2, Pmar-hth1*, and *Pmar hth-2* in situ hybridizations were generated as follows: genes were PCR amplified from cDNA using gene-specific primers designed with Primer3 v. 4.1.0 [101] and complemented with T7 linker sequences (5′ -ggccgcgg-3′; 5′ - cccggggc-3′), and a second PCR using universal T7 primer to generate T7 polymerase templates specific for sense (control) and anti-sense (signal) probes (Additional file 8, Table S2). Sense and antisense riboprobes for *Pmar-Dll* were generated with T7 and T3 polymerases using a plasmid template. For cloning of *Pmar-Dll*, gene-specific primers were used to amplify an 848 bp fragment (Additional file 8, Table S2), which was incorporated to vectors and *E. coli* using TOPO TA cloning kit (Thermo Fisher) following the manufacturer’s instructions, and Sanger sequenced to confirm their identities.

For colorimetric in situ hybridization, we adapted a protocol for the spider *Parasteatoda tepidariorum* after Akiyama-Oda and Oda (2003). The staining reactions lasted between 2h and 8h at room temperature using nitro-blue tetrazolium (NBT) and 5-bromo-4-chloro-3′-indolyphosphate (BCIP) staining reactions (Sigma Aldrich). Embryos were then rinsed in PBS-T (0.05% Tween-20), counter-stained with 10μg/mL Hoechst 33342 (Sigma-Aldrich, St. Louis, MO, USA), post-fixed in 4% formaldehyde/PBS-T and stored at 4°C. Images were taken in a petri dish with agar under PBS-T, using a Nikon SMZ25 fluorescence stereomicroscope mounted with either a DS-Fi2 digital color camera (Nikon Elements software). Appendage mounts were imaged in 70% glycerol/PBS-T in a Zeiss LCM 710 confocal microscope.

## Declarations

### Ethics approval and consent to participate

Studies with arachnids do not required ethics approval or consent to participate.

### Consent for publication

Not applicable.

### Availability of Data and Materials

The data generated and analyzed in this study is included in the main article, supplementary information and available upon request. Raw reads are deposited in NCBI Sequence Read Archive under accession SRR12232018.

### Author’s contributions

Both authors designed the study, analyzed data, developed protocols and contributed to writing. GG conducted gene searches, orthology assessment, phylogenetic analysis, in situ hybridization, assembled figures and drafted the manuscript. Both authors read and approved the final version of the manuscript.

## Acknowledgments

This work is dedicated to Peter Weygoldt, whose pioneer work in the development of *Phrynus marginemaculatus* and contributions to the study of Amblypygi biology continue to inspire the authors of this paper. Comments from two anonymous reviewers improved an early draft of this manuscript. Sequencing was performed at the University of Wisconsin-Madison BioTechnology Center. Microscopy was performed at the Newcomb Imaging Center, Department of Botany, University of Wisconsin-Madison.

## Figure legends

**Additional file 1, Figure S1:**
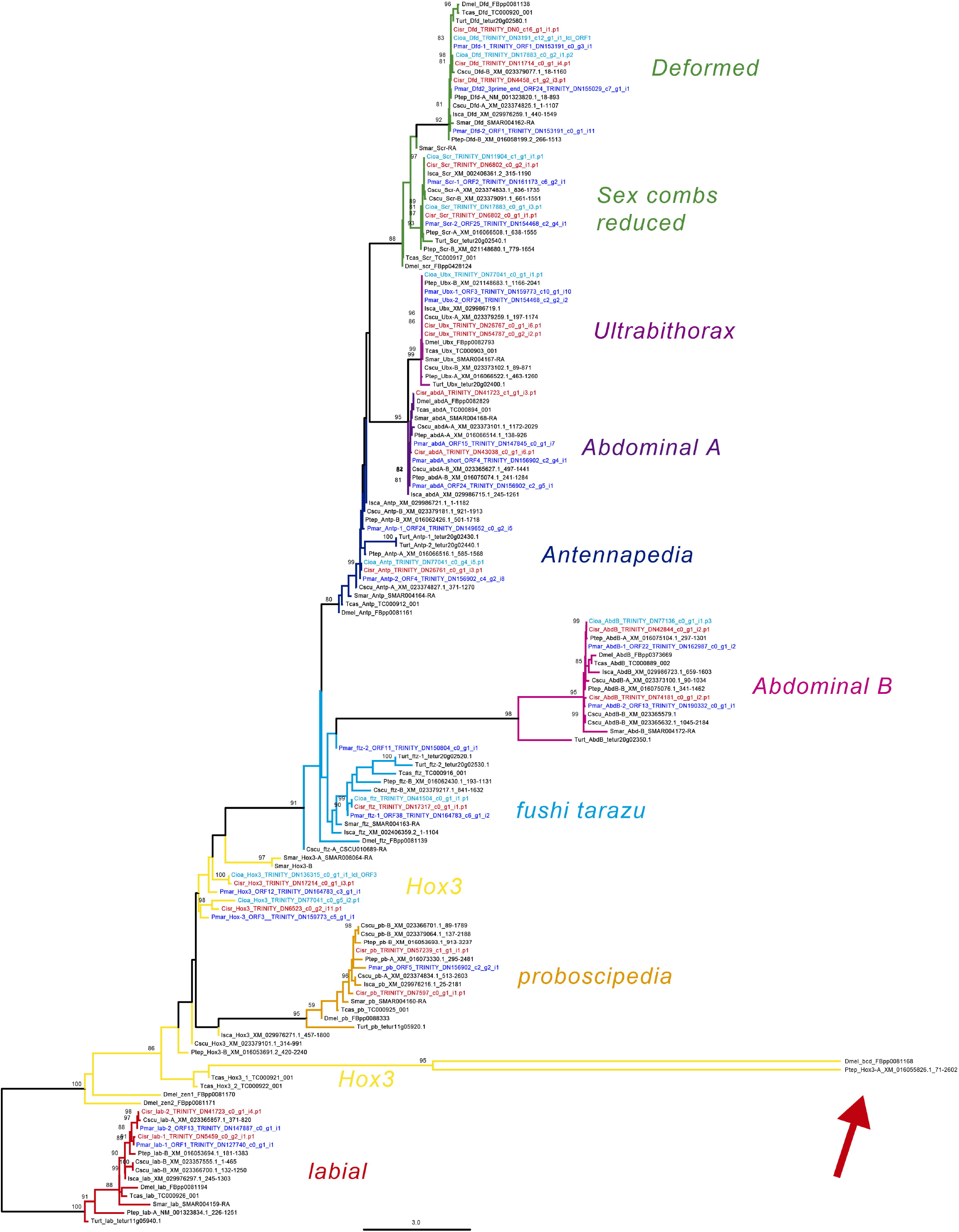
Tree topology inferred from maximum likelihood analysis of a conserved region (75 amino acid characters) (ln L = −4780.167) using the same terminals of Figure 3 and including *Parasteatoda tepidariorum Hox3-A* paralog and *Drosophila melanogaster bicoid* (*Hox3*) (red arrow). Numbers on the notes are ultrafast bootstrap resampling frequencies (only >80 shown). For abbreviations, see Additional file 5, Table S1. File format: jpeg.

**Additional file 2, Figure S2:**
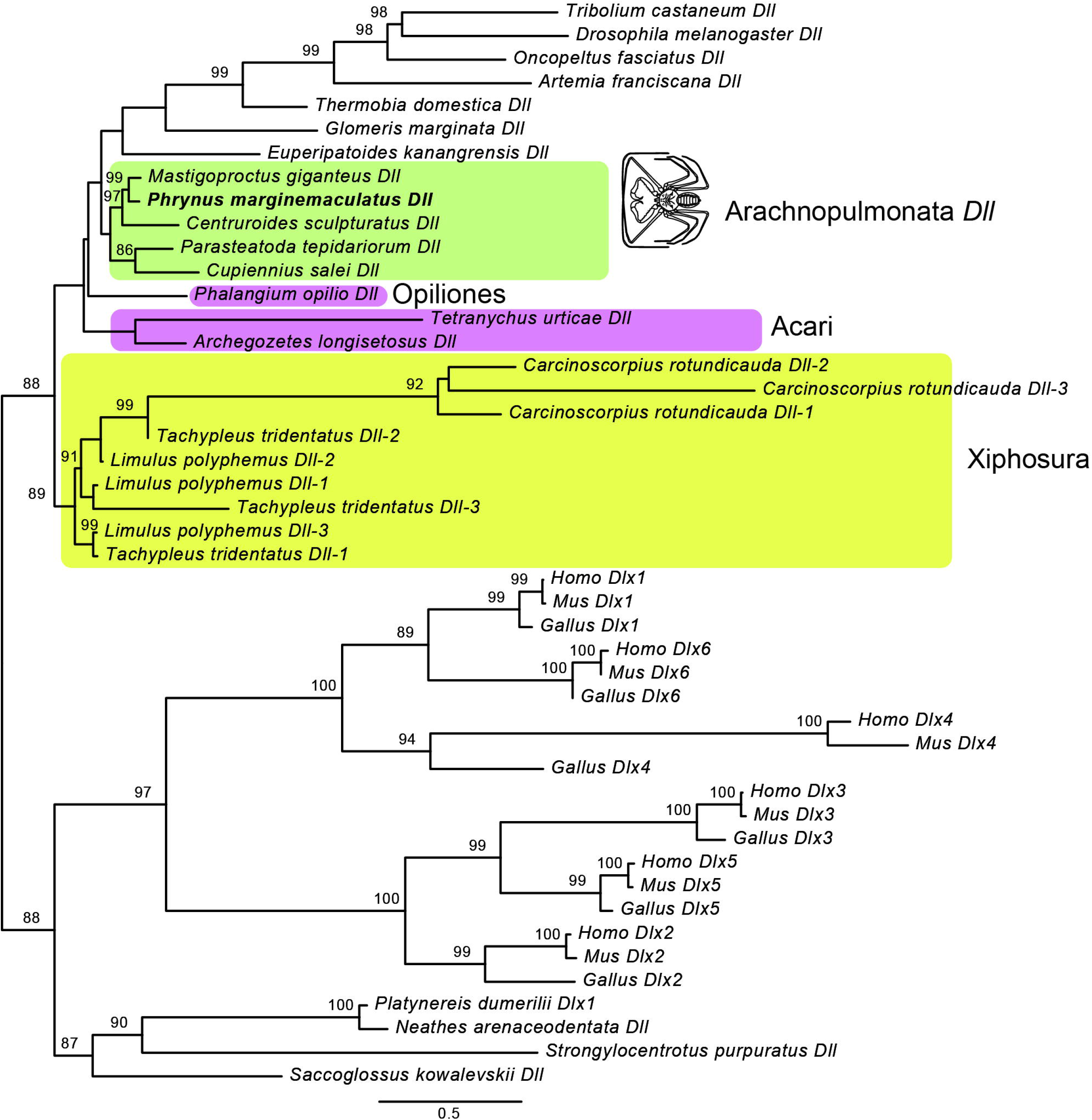
Tree topology of *Distal-less* (*Dll*) inferred from maximum likelihood analysis amino acid sequences (ln L = −15272.780). Numbers on the notes are ultrafast bootstrap resampling frequencies (only >80 shown). Accession numbers are available in Additional file 5, Table S1. The terminal for the whip spider *Phrynus marginemaculatus Dll* ortholog is in boldface. File format: jpeg.

**Additional file 3, Figure S3:**
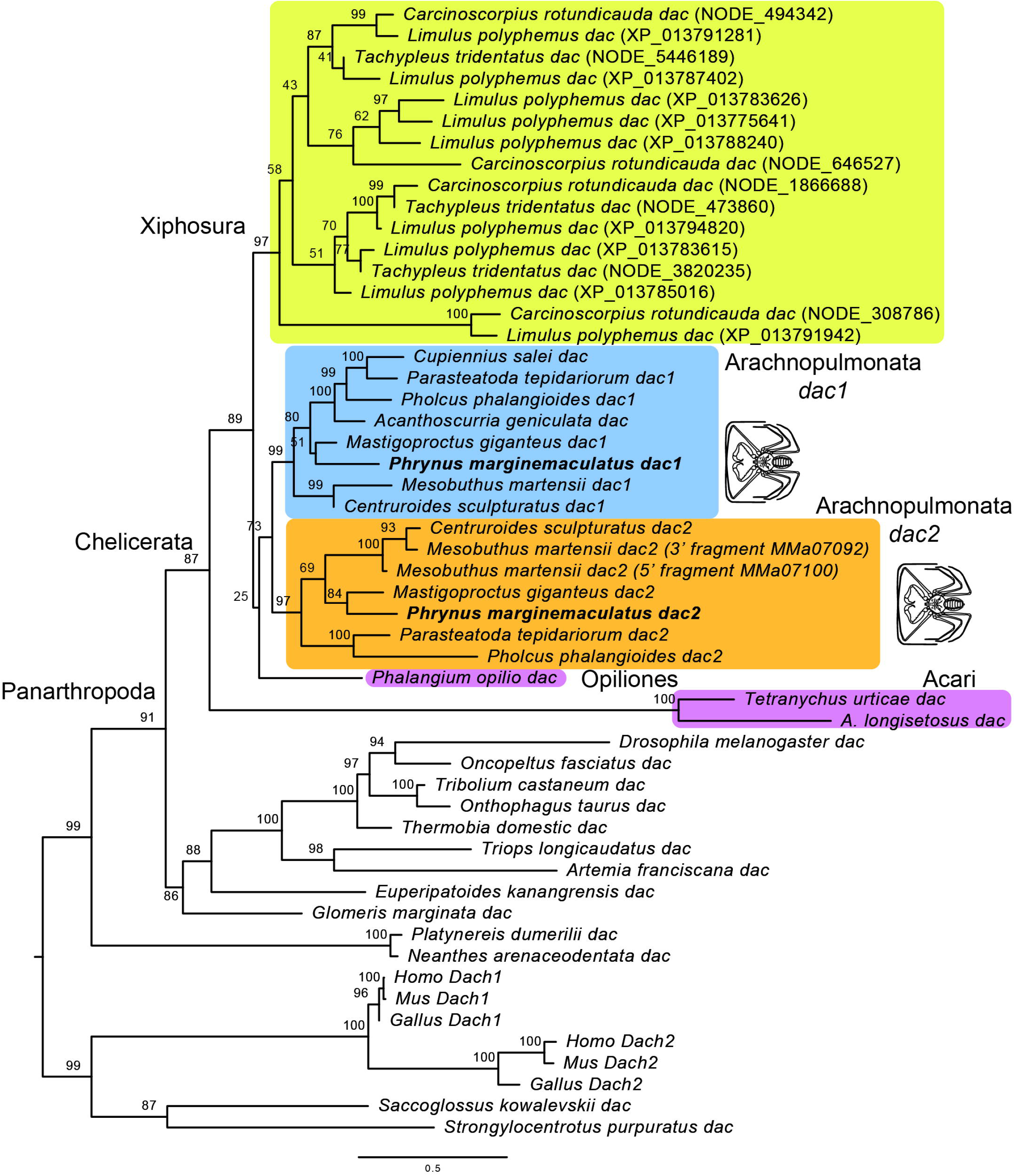
Tree topology of *dachshund* (*dac*) inferred from maximum likelihood analysis amino acid sequences (ln L = −21699.907). Numbers on the notes are ultrafast bootstrap resampling frequencies (only >80 shown). The terminals for the whip spider *Phrynus marginemaculatus Dll* orthologs are in boldface. The original protein alignment of all terminals (except *P. marginemaculatus*) is from Noland et al. 2020. File format: jpeg.

**Additional file 4, Figure S4:**
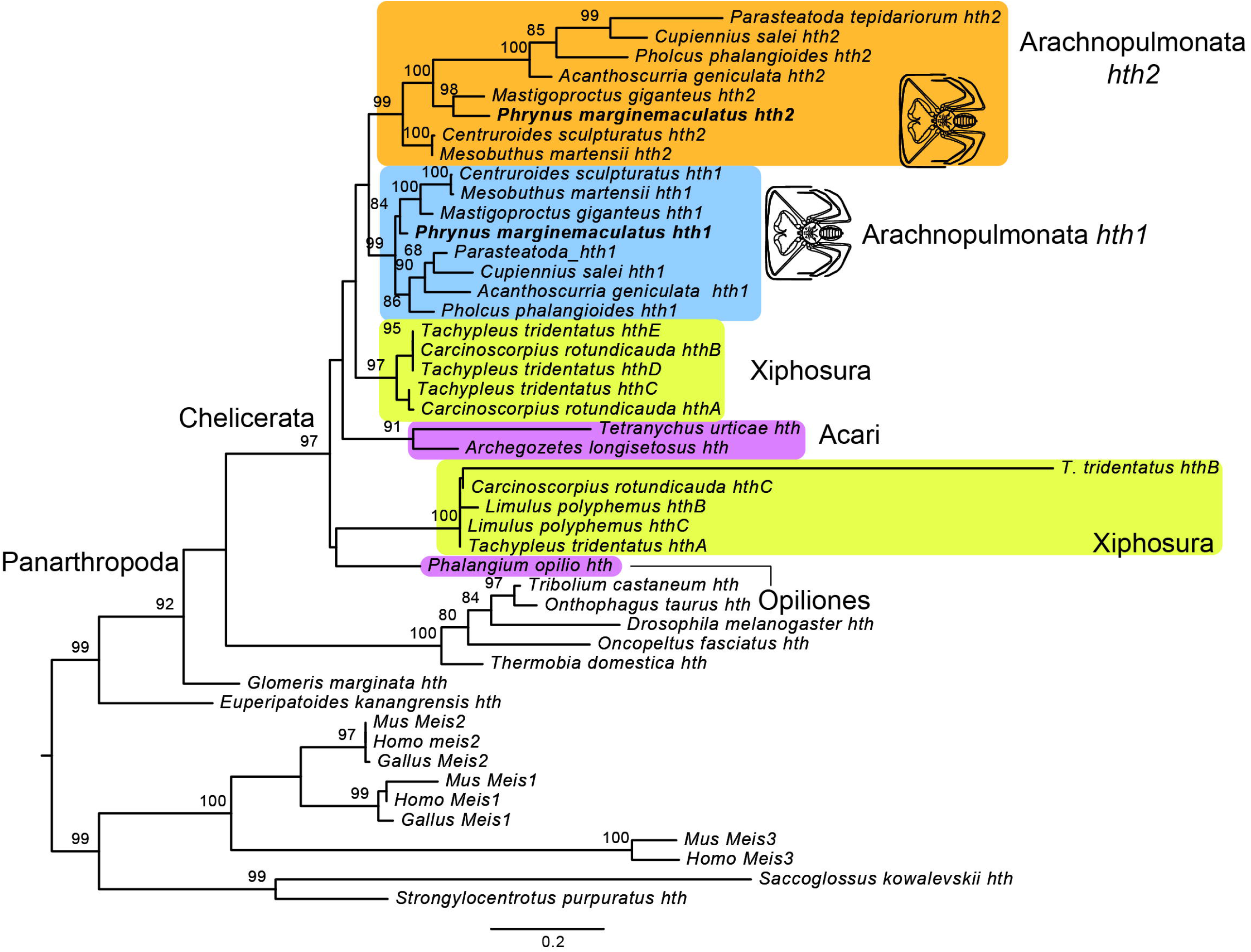
Tree topology of *homothorax* (*hth*) inferred from maximum likelihood analysis amino acid sequences (ln L = −10459.647). Numbers on the notes are ultrafast bootstrap resampling frequencies (only >80 shown). The terminals for the whip spider *Phrynus marginemaculatus hth* orthologs are in boldface. The original protein alignment of all terminals (except *P. marginemaculatus*) is from Noland et al. 2020. File format: jpeg.

Additional file 5, Table S1: List of species, accession numbers and abbreviations of terminals used in the phylogenetic analysis of Hox genes, *Distal-less, dachshund* and *homothorax* homologs. File format: xlsx

Additional file 6: Protein alignment used in the phylogenetic analysis depicted in Figure 3. File format: fasta.

Additional file 7: Protein alignment used in the phylogenetic analysis depicted in Additional file 1, Figure S1. File format: fasta.

Additional file 8, Table S2: List of primers for *Pmar-Dll, Pmar-dac1/2, Pmar hth1/2*, and cloned fragments for *Pmar-Dll*. File format: xlsx

